# Genetic Architecture of Chilling Tolerance in Sorghum Dissected with a Nested Association Mapping Population

**DOI:** 10.1101/622894

**Authors:** Sandeep R. Marla, Gloria Burow, Ratan Chopra, Chad Hayes, Marcus O. Olatoye, Terry Felderhoff, Zhenbin Hu, Rubi Raymundo, Ramasamy Perumal, Geoffrey P. Morris

**Affiliations:** Department of Agronomy, Kansas State University, Manhattan, KS, 66506; USDA-ARS, Plant Stress & Germplasm Development Unit, Cropping Systems Research Lab, Lubbock, TX, 79415; Agricultural Research Center, Kansas State University, Hays, Kansas 67601.

**Keywords:** Multiparental population, Crop evolution, Climate adaptation, Cold tolerance, Antagonistic pleiotropy, Linkage drag

## Abstract

Dissecting the genetic architecture of stress tolerance in crops is critical to understand and improve adaptation. In temperate climates, early planting of chilling-tolerant varieties could provide longer growing seasons and drought escape, but chilling tolerance (<15°) is generally lacking in tropical-origin crops. Here we developed a nested association mapping (NAM) population to dissect the genetic architecture of early-season chilling tolerance in the tropical-origin cereal sorghum *(Sorghum bicolor* [L.] Moench). The NAM resource, developed from reference line BTx623 and three chilling-tolerant Chinese lines, is comprised of 771 recombinant inbred lines genotyped by sequencing at 43,320 single nucleotide polymorphisms. We phenotyped the NAM population for emergence, seedling vigor, and agronomic traits (>75,000 data points from ∼16,000 plots) in multi-environment field trials in Kansas under natural chilling stress (sown 30–45 days early) and normal growing conditions. Joint linkage mapping with early-planted field phenotypes revealed an oligogenic architecture, with 5–10 chilling tolerance loci explaining 20–41% of variation. Surprisingly, several of the major chilling tolerance loci co-localize precisely with the classical grain tannin (*Tan1* and *Tan2*) and dwarfing genes (*Dw1* and *Dw3*) that were under strong directional selection in the US during the 20th century. These findings suggest that chilling sensitivity was inadvertently selected due to coinheritance with desired nontannin and dwarfing alleles. The characterization of genetic architecture with NAM reveals why past chilling tolerance breeding was stymied and provides a path for genomics-enabled breeding of chilling tolerance.

**Article Summary:** Chilling sensitivity limits productivity of tropical-origin crops in temperate climates, and remains poorly understood at a genetic level. We developed a nested association mapping resource in sorghum, a tropical-origin cereal, to understand the genetic architecture of chilling tolerance. Linkage mapping of growth traits from early-planted field trials revealed several major chilling tolerance loci, including some colocalized with genes that were selected in the origin of US grain sorghum. These findings suggest chilling sensitivity was inadvertently selected during 20th century breeding, but can be bypassed using a better understanding of the underlying genetic architecture.

**Disclaimer:** Mention of a trademark, warranty, proprietary product, or vendor does not constitute a guarantee by the USDA and does not imply approval or recommendation of the product to the exclusion of others that may be suitable. USDA is an equal opportunity provider and employer.

## Introduction

Adaptation to diverse environments has generated abundant genetic diversity in wild and domesticated plant species (Anderson *et al.* 2011; Meyer and Purugganan 2013). The genetic architecture of adaptation has been intensively studied both theoretically and empirically, but remains contentious. For instance, much debate surrounds the relative contributions of standing genetic variation versus new mutation (Barrett and Schluter 2008), oligogenic versus polygenic variation (Orr 2005), and pleiotropic versus independent effects (Paaby and Rockman 2013). Despite the importance of adaptive variation in crop improvement, the genomic basis of local adaptation underlying abiotic stressors remains poorly understood (Olsen and Wendel 2013). Understanding the genomic basis of adaptation in crops can guide breeding strategies and facilitate transfer of adaptive traits for new climate-resilient varieties (Soyk *et al.* 2017; Zhu *et al.* 2018; Li *et al.* 2018).

Cold temperatures are a major factor limiting plant productivity globally for both wild plants and crops (Cramer *et al.* 1999). Tropical-origin crops (e.g. maize, rice, tomato, cotton, sorghum) are typically sensitive to chilling temperatures (0–15°), which limits their range and/or growing season in temperate climates (Lyons 1973; Long and Spence 2013). Developing chilling-tolerant varieties could facilitate early planting to extend growing seasons, prevent soil moisture depletion, and shift growth and flowering to more favorable evapotranspirative conditions (Tuberosa 2012; Ma *et al.* 2015). For breeding chilling tolerance in tropical-origin crops, chilling-adapted germplasm from high-latitude zones and high-altitude tropical regions can be targeted as donors. Molecular mechanisms underlying cold tolerance (chilling and/or freezing temperatures) include C-repeat binding factor (CBF) regulon cold signaling (Thomashow 2001; Park *et al.* 2015; Wang *et al.* 2018), jasmonate signaling (Hu *et al.* 2013; Mao *et al.* 2019), and lipid remodelling (Li *et al.* 2004; Moellering *et al.* 2010).

Sorghum, a tropical-origin warm-season (C4) cereal, is among the major crops that are generally susceptible to chilling (Franks *et al.* 2006). Sorghum originated in tropical Africa (c. 5–10 thousand years ago) and diffused to temperate areas, including China (c. 800 years ago) and the United States (c. 200 years ago) (Kimber 2000). Diffusion of tropical sorghums to temperate climates has led to commercial sorghum industries covering several million hectares in US, Australia, Argentina, and China (Monk *et al.* 2014). Using a phytogeographic approach (Vavilov 1951), Chinese sorghum were targeted as chilling tolerance donors for conventional breeding starting in the 1960s (Stickler *et al.* 1962). However, characteristics of Chinese sorghums that are undesirable for US grain sorghum, particularly grain tannins and tall stature (>2 m) (Franks *et al.* 2006), have hampered breeding. Biparental linkage mapping identified chilling tolerance QTL tagged by the same molecular markers as grain tannins and plant height, but small populations and low marker density limited dissection of these traits (Knoll *et al.* 2008; Brown *et al.* 2008; Xiang 2009; Burow *et al.* 2010; Wu *et al.* 2012). Several classical tannin (*Tan1* and *Tan2*) and dwarfing (*Dw1*–*Dw4*) genes (Stephens 1946; Quinby and Karper 1954) have been cloned in recent years (Multani *et al.* 2003; Wu *et al.* 2012; Hilley *et al.* 2016, 2017), which could aid further trait dissection.

NAM populations can provide increased power for dissecting complex traits (Buckler *et al.* 2009; Ogut *et al.* 2015), particularly for adaptive traits where population structure confounds associations studies of natural populations (Bouchet *et al.* 2017). To dissect the genetic architecture of early-season chilling tolerance in sorghum, we developed and deployed a new nested association mapping (chilling NAM) resource. The chilling NAM population addresses a gap in existing sorghum NAM resources (Bouchet *et al.* 2017) by including contrasting temperate-adapted founders, three chilling-tolerant Chinese founders and the chilling-susceptible reference line BTx623 (Paterson *et al.* 2009). We used the chilling NAM to dissect the genetic architecture of sorghum early-season chilling tolerance at a high resolution based on natural field stress conditions. This NAM study provides insights into the origin and persistence of chilling sensitivity in US grain sorghum, and reveals new strategies for genomics-enabled breeding in this system.

## Materials and Methods

### Population development

The chilling NAM population consists of three biparental populations that share a common parent, the US reference line BTx623 (Paterson *et al.* 2009) (Figure S1). The NAM founders were selected based on their contrasting chilling responses from early planting in preliminary studies in Lubbock, Texas. Chilling-sensitive BTx623 was used as the seed parent in crosses with three chilling-tolerant Chinese founders, Niu Sheng Zui (NSZ; PI 568016), Hong Ke Zi (HKZ; PI 567946), and Kaoliang (Kao; PI 562744) in Lubbock, Texas. BTx623 is derived from Combine Kafir × SC170, an Ethiopian zerazera caudatum (Menz *et al.* 2004). The resulting F_1_ progenies were self-pollinated to generate three segregating F_2_ populations. RILs were developed using single-seed descent by selfing to the F_6_ generation in Lubbock, Texas (summer nursery) and Guayanilla, Puerto Rico (winter nursery). The F_6:7_ RILs were derived by combining seeds of 3–4 uniform panicles. Additional seed increase of the NAM population was conducted in Puerto Vallarta, Mexico (winter nursery), by selfing the F_6:7_ plants to derive F_6:8_ RILs. Below, the Chinese founder name will be used when referring to a given RIL family (e.g., the NSZ family).

### Early-and normal-planted field trials

Six early-and two normal-planted field trials were conducted in 2016, 2017, and 2018 in Kansas (Table S1). Three locations, two in eastern Kansas [Ashland Bottoms (AB), 39.14N -96.63W; Manhattan (MN), 39.21N -96.60W] and one in western Kansas [Agricultural Research Center, Hays (HA), 38.86N -99.33W], were used for field trials (Figure 1A). Abbreviated location name and the last two digits of the year (e.g. AB16 for Ashland Bottoms 2016) were assigned for each field trial. A suffix was added to the AB16 field trials, AB16_b1 and AB16_b2, as both were planted in AB with an interval of two weeks between plantings. The F_6:7_ RILs were planted in AB16, while F_6:8_ RILs were planted in AB, MN, and HA in 2017 and 2018. Each field trial contained two replicates of the NAM population. The NAM RILs were randomized within family in 2016, and completely randomized in 2017 and 2018 in each replicate block (Figures 1A and S2). Controls in each field trial comprised chilling-tolerant Chinese accessions NSZ, HKZ, Kao, and Shan Qui Red (SQR; PI 656025), chilling-sensitive inbreds BTx623 and RTx430, and US commercial grain sorghum hybrid Pioneer 84G62.

**Figure 1.**
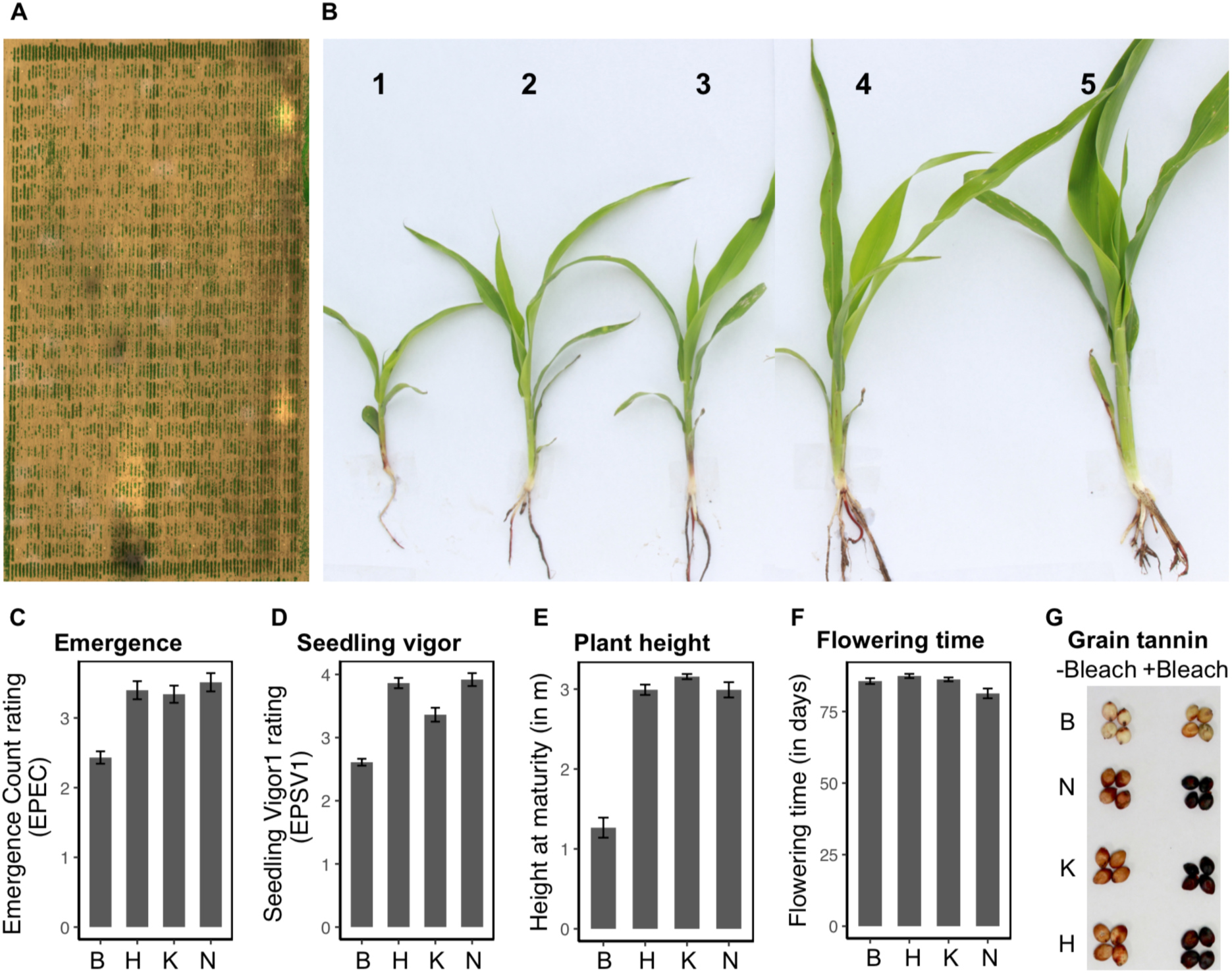
Chinese sorghums harbor early-season chilling tolerance and characteristics undesirable for US grain sorghums. (A) Aerial image of an early-planted field (AB17) trial for chilling tolerance phenotyping based on stitched RGB imagery (B) Seedling vigor rating used in field trials. In early-planted field trials, differences were observed in (C) emergence and (D) seedling vigor between the four NAM founders, B (BTx623), K (Kao), H (HKZ), and N (NSZ). Additionally, (E) significant variation in plant height at maturity, (F) no significant difference in flowering time (days after emergence), and (G) presence/absence variation in grain tannins were observed.

Five early-planted (EP, natural chilling stress) trials were sown in April and one in early May (MN17), 30–45 days earlier than normal sorghum planting in Kansas (Grain Sorghum Production Handbook, 1998). The EP trials, except MN17, experienced chilling stress (<15°) during emergence (Table S1 and Figure S3). Optimal temperatures (>15°) prevailed in MN17 during emergence, but one-week-old seedlings experienced chilling stress (5–13°). Normal-planted (NP, optimal temperature) field trial was sown in June when the soil temperatures were optimal for sorghum cultivation (>15°). AB18 was considered as the second NP trial, although planted in early May, as optimal conditions prevailed during emergence and seedling growth.

### Field phenotyping

Seedling phenotypes of the NAM population were evaluated under early-and normal-planted field trials. Prefixes EP and NP were included for each seedling trait to differentiate phenotypes from early-and normal-planted trials, respectively. Emergence count (EC) was scored on a scale of 1–5 that represented 20, 40, 60, 80, and 100% emergence, respectively. Three seedling vigor (SV) ratings (SV1–SV3) were collected at week-1, -2, and -4, respectively, after emergence. SV was scored on a rating scale of 1–5 with a rating of 1 and 5 for low and robust vigor, respectively (Figure 1B). A previously described SV scale (Maiti *et al.* 1981) was modified (1 for high and 5 for low SV) for consistency with EC rating. Repeatability of SV rating, SV2 (AB17) and SV3 (MN17), was tested with SV ratings collected simultaneously by different individuals. Early-planted damage rating (EPDR), based on visual damage observed two days after a severe chilling stress event, was scored on a 1–5 rating scale representing seedling death/severe leaf-tip burning, leaf-tip burning, severe chlorosis, mild/partial chlorosis, and no chilling damage symptoms, respectively. Seedling height was measured manually one month after initial emergence in each location.

Plant height and flowering time (days to flowering after emergence), the major agronomic traits, were collected from three (AB16_b1, MN17, and MN18) and two (AB16_b1 and MN17) field trials, respectively. Agronomic suitability of the NAM population as US grain sorghum, which included semi-dwarf stature, panicle exertion, standability, and compact panicle architecture, was screened in AB16_b1. Presence or absence of grain tannins in field-grown samples (from Puerto Vallarta, Mexico) of each RIL was determined using bleach test with SQR, a Chinese accession containing grain tannin, as a positive control (Wu *et al.* 2012). Fifteen seeds from each RIL were transferred into a 2 ml tube and 1 ml bleach solution (3.5% sodium hypochlorite and 5% sodium hydroxide) was added. RILs with tannins turned black after 30 min and were scored as 1. By contrast, nontannin RIL seed did not change their color and were scored as 0.

### Statistical analysis of phenotypes

Trait correlation between locations was determined using the averaged seedling trait ratings of two replicates from each field trial. Pearson pairwise correlation analysis was performed using *pairs.panels* function in psych R package. Broad sense heritability (*H*^2^) estimate of EP and NP field phenotypes was calculated with seedling ratings from six EP and two NP field trials, respectively. Seedling traits *H*^2^ was calculated from variance components generated with the *lme4* (Bates *et al.* 2015) R package as described earlier (Boyles *et al.* 2017). All components were treated as random effects and replicates were nested in location-by-year interaction:

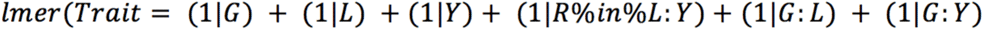

and broad-sense heritability was calculated using the equation:

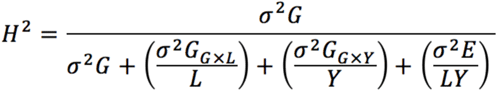

where *G* is the genotype, *L* is the location, *Y* is the year, *R* is the replicate, and *E* is the error term. Environment main effects were not included in the denominator as they do not influence response to selection (Holland *et al.* 2003). Best linear unbiased predictions (BLUPs) of EP and NP seedling traits were generated using the model for estimating *H*^2^.

### DNA extraction and genotyping

Genotyping-by-sequencing (GBS) was conducted on one-week-old seedlings of the F_6:7_ RILs and the four NAM founders (Figure S1). Leaf tissue (∼50 mg, pooled from three seedlings) from each RIL was transferred into a 96-deepwell plate, lyophilized, and stored at -80°. One ball bearing was added to each well and the leaf tissue was ground with a Retsch Mixer Mill MM400 tissue grinder (Vernon Hills, IL, USA). Genomic DNA was extracted using QIAGEN BioSprint 96 DNA plant kit (Germantown, MD, USA). DNA was quantified with Quant-iT PicoGreen dsDNA assay kit (Thermo Fisher Scientific, Grand Island, NY, USA) using Agilent 2100 Bioanalyzer (Santa Clara, CA, USA) at the Kansas State University Integrated Genomics Facility. Each sample was normalized to contain 10 ng/µl DNA using QIAgility Liquid Handling System (Germantown, MD, USA). Six µl of DNA was transferred to a 96-well PCR plate and adapters were added. *Ape*KI enzyme was used for restriction digestion and GBS libraries were prepared as described previously (Elshire *et al.* 2011; Morris *et al.* 2013a). Illumina HiSeq 2500 Rapid v2 sequencing system was used for 100-cycle single-end sequencing of two 384-multiplexed libraries at the University of Kansas Medical Center Genome Sequencing Facility.

GBS data from the chilling NAM resource was combined with previously published *Ape*KI GBS data from ∼10,323 diverse accessions (Hu *et al.* 2019), aligned to the BTx623 reference genome v3.1 (McCormick *et al.* 2018), and SNP calling was performed using Tassel 5.0 GBS v2 pipeline (Glaubitz *et al.* 2014). GBS of the chilling NAM population provided 528,065 single nucleotide polymorphisms (SNPs) (Figure S1). After filtering the GBS data for 80 percent missingness (PM) and 0.05 minor allele frequency (MAF) 61,428 SNPs were retained. These SNPs were separated by individual chromosomes and imputed using Beagle 4.1 (Browning and Browning 2013). Additional filtering for markers and RILs with >15% residual heterozygosity retained 43,320 SNPs and 750 RILs for joint linkage mapping.

### Population genetic analyses

Genetic structure of the chilling NAM population was characterized with respect to global sorghum germplasm. First, the chilling NAM and global accessions GBS data was filtered for 80 PM and 0.01 MAF, and the retained SNPs (265K) were imputed using Beagle 4.1 (Browning and Browning 2013). Next, two PCA axes were built with previously published *Ape*KI GBS data of 401 global sorghum accessions (Morris *et al.* 2013a), and chilling NAM founders and RILs were projected on these axes. Principal component analysis (PCA) of global germplasm was performed using *prcomp* function in R. Coordinates for the chilling NAM population were calculated with the *predict* function in R.

Neighbor-joining analysis, using TASSEL 5.0, was conducted with 61,428 SNPs to characterize the genetic relatedness of the chilling NAM population. Phylogenetic tree was constructed with Ape package (Paradis *et al*. 2004) in R (R Core Team, 2014). SNP density was calculated with VCFtools in 200kb windows. Linkage disequilibrium (LD) decay was estimated, using pairwise comparisons of ∼55–70K GBS SNPs, individually for the three NAM families with PopLDdecay v.3.29 package (Zhang *et al.* 2018). LD decay of 176 Ethiopian and 29 Chinese landraces (genotyped previously with *Ape*KI) (Lasky *et al.* 2015) was estimated for comparison. Ethiopian and Chinese germplasm LD decay was calculated using ∼100K and ∼57K SNPs, respectively. Parameters were set for -MaxDist as 500 kb and -MAF as 0.05. LD decay curves were plotted based on *r*^*2*^ and the distance between pairs of SNPs.

### Linkage mapping analysis

The NAM founders genotypes were used for constructing genetic linkage maps with the R/qtl package (Broman *et al.* 2003). The NAM founders were filtered for 20 PM and <0.4 MAF and the retained SNPs were used to retrieve the NAM population genotypes from the GBS dataset. SNP imputation was conducted for each family separately using Beagle 4.1 (Browning and Browning 2016). RILs with >85% missing data or >80 crossovers were dropped. Duplicate markers (i.e. mapping to the same location) were dropped. Genetic linkage maps for each NAM family were generated using the *Haldane* function. The *Droponemarker* function in R/qtl was used to discard problematic markers that increase chromosome length. Genetic linkage maps were reconstructed for each NAM family. Composite interval mapping (CIM) (Zeng 1994), with R/qtl, was used for performing linkage mapping and significant QTL were determined based on the threshold level defined by computing 1000 permutations. Allelic effects were defined as positive or negative effects of the BTx623 allele. LOD support interval for individual QTL was obtained with the *lodint* R/qtl function. CIM was performed with plant height, flowering time, and grain tannin data to validate the generated genetic linkage maps. BLUPs of seedling traits, EC and SV1–3, from early-and normal-planted field trials were used for CIM. Additionally, linkage mapping was performed for individual field trials with the averaged data of two replicates from each location.

### Joint linkage mapping

Joint linkage mapping (JLM) was conducted with 43,320 GBS SNPs and seedling trait BLUPs from 750 RILs. In addition, JLM was performed individually for each location with the averaged sdata of two replicates. Mapping power and resolution of the chilling NAM population was validated using plant height, flowering time, and grain tannin data. Stepwise regression approach in TASSEL 5.0 (Glaubitz *et al.* 2014), which uses forward inclusion and backward elimination stepwise method, was used to perform JLM. Entry and exit limit of the forward and backward stepwise regressions was 0.001 and threshold cut off was set based on 1000 permutations. JLM was performed using the following equation:

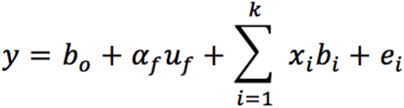

where b_0_ is the intercept, *u*_*f*_ is the effect of the family of founder line f obtained in the cross with the common parent (BTx623), *α*_*f*_ is the coefficient matrix relating *u*_*f*_ to y, *b*_*i*_ is the effect of the ith identified locus in the model, *x*_*i*_ is the incidence vector that relates *b*_*i*_ to y and k is the number of significant QTL in the final model. Allelic effect for each QTL was expressed relative to the BTx623 allele, where alleles with positive-or negative-additive effects were derived from BTx623 or Chinese founders, respectively. Based on the average genome-wide recombination rate of 2.0 cM/Mb for sorghum (Mace *et al.* 2009; Bouchet *et al.* 2017), QTL for one or more seedling traits that mapped within a 2 Mb interval were assigned a common name. For example, *qSbCT04.62* to describe QTL detected on chromosome 4 close to 62 Mb.

### Sequence variant analysis

*CBF* and *Tan1* genes, colocalizing with chilling tolerance QTL, were used for sequence variant analysis. Two overlapping primer pairs were used to amplify these genes from the Chinese founders (primer sequences are included in Table S2). 50% glycerol and 25mM MgCl_2_ were added to the master mix for stabilizing the PCR reaction. PCR product purification and Sanger sequencing were performed at GENEWIZ (South Plainfield, NJ). Clustal Omega and Expasy translate were used for sequence alignment and predicting the peptide sequences of *CBF1* and *Tan1* genes.

### Ecophysiological crop modeling

CERES-Sorghum crop model (White *et al.* 2015) in the Decision Support Systems for Agro-technology Transfer-Crop Simulation Model software (Jones *et al.* 2003; Hoogenboom *et al.* 2017) was used to predict the value of early planting for grain sorghum in the Kansas production environment. This model simulates daily physiological processes using a base temperature of 8° (White *et al.* 2015) and has effectively predicted sorghum grain yield in Kansas (Staggenborg and Vanderlip 2005; Araya *et al.* 2018). We consider that this model assumes chilling tolerance by default, since it does not model damage due to chilling temperatures. A full-season (late-maturing) photoperiod insensitive grain sorghum hybrid, used in previous crop modelling, was used in this study (Araya *et al.* 2018). Simulations were performed under rainfed conditions at four representative Kansas locations, Colby (39.39N, -101.06W), Garden City (37.99N, -101.81W), Hays (38.84N, -99.34W), and Manhattan (39.20N, -96.55W), from a 30 year period (1986 to 2015). Historical weather data for each of these locations was obtained from Kansas Mesonet (2019). Simulations were started on January 1 to account for the effect of precipitation on soil moisture and the onset of soil evaporation. Early (April 15), normal (May 15), and late (June 15) planting scenarios were simulated, and (i) available precipitation, (ii) days of water stress after anthesis, and (iii) final grain yield were analyzed.

### Data availability

Sequencing data are available in the NCBI Sequence Read Archive under project accession SRP8838986. Field phenotyping data and R analysis scripts are deposited in Dryad Digital Repository (doi: *[Add after acceptance]*). Plant materials: The chilling NAM population seeds will be submitted to the USDA National Plant Germplasm System’s Germplasm Resource Information Network (https://www.ars-grin.gov/). Please contact G.B. or the corresponding author for availability.

## Results

### Development of NAM population for chilling tolerance studies

The chilling NAM population was generated from crosses of a US reference line BTx623 with three Chinese lines, NSZ, Kao, and HKZ (Figure S1). The resulting chilling NAM population (*n* = 771) comprised 293, 256, and 222 RILs for the NSZ, Kao, and HKZ families, respectively. Our chilling tolerance studies of the NAM founders and RILs were based on natural chilling events in field trials sown 30-45 days earlier than normal. In early-planted field trial (Figures 1A–B) the Chinese founders had significantly greater emergence and seedling vigor (*P* < 0.05) than BTx623 (Figures 1C, 1D, and S4). Chinese founder lines were much taller (∼3 m) at maturity than BTx623 (1.2 m) (Figure 1E), but little variation was observed for flowering time among the founder lines (4–5 d; *P* < 0.05; Figure 1F). Grain tannins were present in the Chinese accessions and absent in BTx623 (Figure 1G).

### Genetic properties of the chilling NAM population

The filtered GBS data set for the chilling NAM population comprised genotypes at 43,320 SNPs. SNP densities were higher in telomeres than pericentromeric regions (Figure S5A). To check the population’s quality and understand its genetic structure, NAM RILs and founders were projected onto PCA axes built from a global sorghum diversity panel (Figure 2A), which reflect geographic origin and botanical race (Harlan *et al.* 1972; Morris *et al.* 2013a). As expected, the Chinese founders clustered with durra sorghums of Asia and East Africa, while BTx623 was positioned midway between kafir and caudatum clusters, consistent with its pedigree (Menz *et al.* 2004) (Figure 2A). The three half-sib families of the chilling NAM population were clustered together, midway between the Chinese founders and BTx623. NJ analysis (Figure S5B) and PCA (Figure S5C) of the chilling NAM population by itself confirmed the expected family structure for NSZ and Kao, with each family forming a single cluster. Two clusters were observed for the HKZ family. We assigned HKZ RILs into HKZa (n_RIL_ = 121) or HKZb (n_RIL_ = 101) subfamilies, with the HKZb subfamily representing the cluster with PC1 > 40 (and the longer branch on NJ dendrogram). The LD rate decay (to genome-wide background) was slower in NAM families (∼500 kb) compared to diverse accessions from China and Ethiopia (∼20 kb) (Figure 2B).

**Figure 2.**
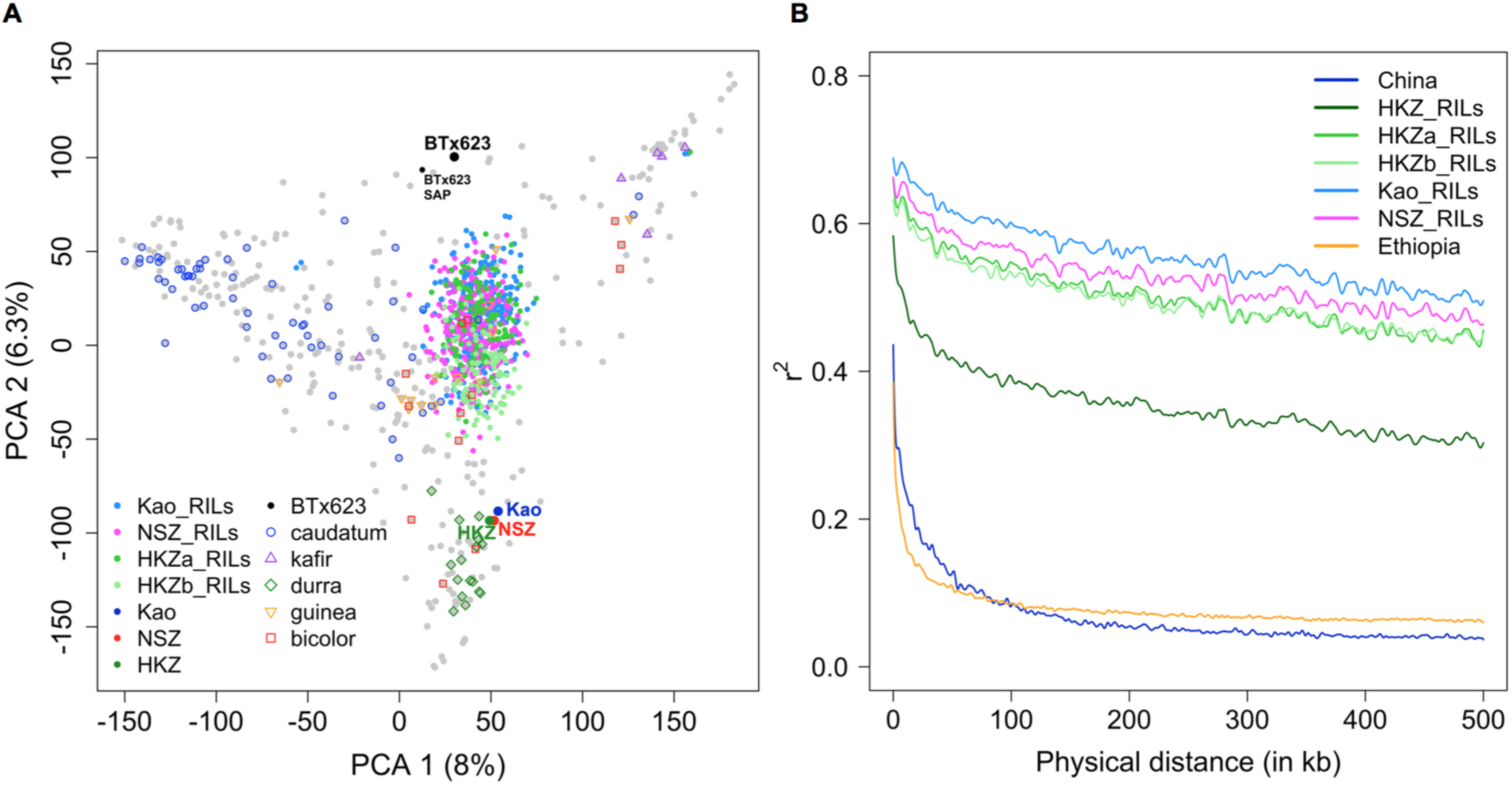
Genetic properties of the chilling NAM population. (A) Principal component analysis (PCA) of the NAM (n_RIL_ = 771) plotted on PCA axes built with 401 accessions of the global sorghum diversity germplasm. Major botanical races (Caudatum, Kafir, Durra, Guinea, and Bicolor) of global accessions are noted with symbols (B) Linkage disequilibrium (LD) decay of the NSZ, HKZa, HKZb, and Kao families. LD decay rate of diverse accessions from China (n = 29) and Ethiopia (n = 176) are presented for comparison.

### Repeatability and heritability of field phenotypes

RILs were scored for emergence and seedling vigor under early-and normal-planted field trials. Early (EPSV1) and later (EPSV2, EPSV3) seedling vigor ratings were strongly correlated (0.7– 0.8), as were ratings made by different individuals on the same day (0.7–0.8) (Figure S6). By contrast, the correlation across RILs between early-and normal-planted seedling traits was low (0.1–0.3). Broad sense heritability (*H*^2^) across locations and years for early-planted seedling traits was intermediate (0.4–0.5) (Table 1), while *H*^2^ was higher (0.5–0.8) for seedling traits from normal-planted field trials. *H*^2^ for seedling height (in early-planted field trials) was close to zero (0.03), while plant height at maturity was highly heritable (0.9). Based on the averaged data of two replicates within each field trial, low to intermediate correlation (0.1–0.4) was observed with the same seedling trait among locations for early-planted trials (Figure S7).

**Table 1.** Broad-sense heritability (*H*^*2*^) of early-and normal-planted field traits.

| Seedling traits | $H^2$ early planting<sup>a</sup> | $H^2$ normal planting<sup>b</sup> |
| --- | --- | --- |
| Emergence count (EC) | 0.45 | 0.53 |
| Seedling vigor1 (SV1) | 0.52 | 0.57 |
| Seedling vigor2 (SV2) | 0.39 | 0.78 |
| Seedling vigor3 (SV3) | 0.37 | 0.53 |
| Damage rating (DR) | 0.35 | - |
| Seedling height | 0.03 | - |
| Plant height at maturity | 0.93 | - |

<sup>a</sup> Field phenotypes from six early-planted trials.
<sup>b</sup> Field phenotypes from two normal-planted trials.

### Composite interval mapping of early-season chilling tolerance

Genetic linkage maps were constructed for each family (NSZ: 1341 markers, 257 RILs; Kao: 1043 markers, 219 RILs; HKZa: 1150 markers, 107 RILs) (Figure S8). Map lengths were similar for the NSZ, Kao, and HKZa families (1403 cM, 1381 cM, and 1295 cM, respectively) and individual RILs contained 2–4 crossovers. To map putative chilling tolerance loci, composite interval mapping (CIM) was first conducted in individual families using ∼1000–1300 markers and early-planted seedling trait BLUPs (EPEC, EPSV1–3). CIM detected 6–8 QTL, which explained 16–28%, 8–23%, and 12–36% of variation for early-planted seedling traits in the HKZa, Kao, and NSZ families, respectively (Table S3). The QTL on chromosome 4 was detected in all NAM families, with the positive allele inherited from the Chinese founder in each case. CIM of normal-planted seedling BLUPs (NPEC and NPSV1–NPSV3) identified 4–9 QTL contributing to emergence and SV in the HKZa, Kao, and NSZ families, respectively. Few overlaps were observed among QTL detected for early-and normal-planted seedling traits (Tables S3 and S4). As chilling stress varied among locations (Figure S3), QTL mapping was conducted for each field trial separately to check the stability of QTL across locations. The QTL on chromosomes 4 and 7 were detected across families in four and two early-planted trials, respectively.

### Joint linkage mapping of early-season chilling tolerance

To leverage data across families, JLM was performed with 43,320 SNPs and field phenotypes from 750 RILs (including the HKZb family) (Figure 3A–E). JLM of seedling trait BLUPs (derived from ∼12,000 early-planted plots) identified 15 QTL, seven of which were detected for multiple seedling traits (Figure 3D and Table 2). Each QTL explained 1–9% of phenotypic variation. In total, the QTL explained 21–41% variation for emergence and seedling vigor. Positive alleles were inherited from the Chinese founders, except for the allele at chromosome 3. The QTL on chromosomes 2 and 4 were detected for every early-planted seedling trait. The chromosome 1 and 5 QTL were detected with all seedling vigor traits, while chromosome 7 and 9 were mapped with two early-planted seedling traits (Figure 3D). The QTL on chromosomes 2 and 4 colocalized (<1 Mb) with classical tannin genes, *Tan2* and *Tan1* (Wu *et al.* 2012; Morris *et al.* 2013b), and chromosomes 7 and 9 loci colocalized with classical dwarfing genes, *Dw3* and *Dw1* (Multani *et al.* 2003; Hilley *et al.* 2016). JLM of normal-planted traits mapped different QTL for emergence, but few overlapped with QTL for early-planted seedling vigor (Figures 3C and S9–S12, and Table S5).

**Table 2.**
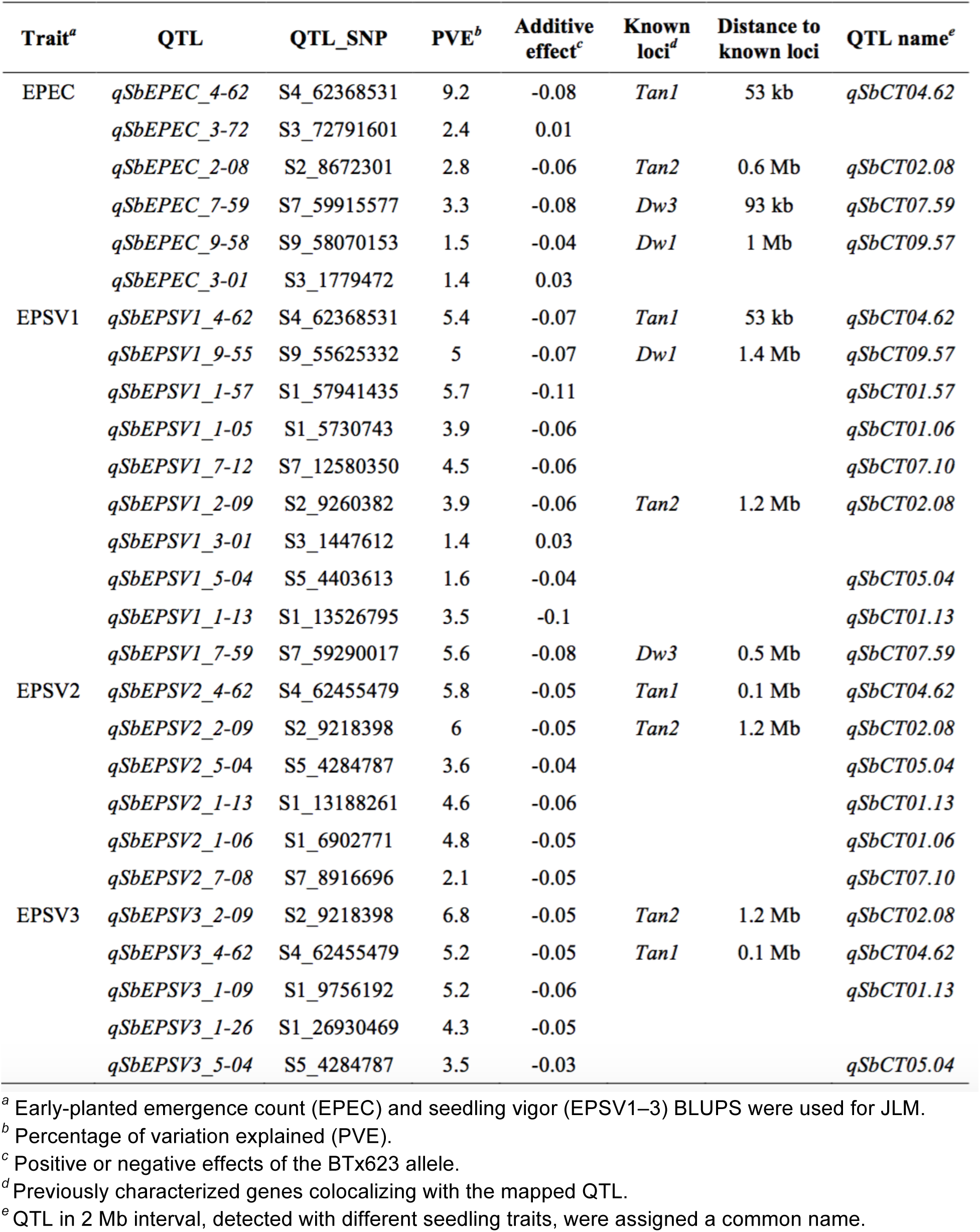
Joint linkage mapping (JLM) with early-planted field phenotypes.

**Figure 3.**
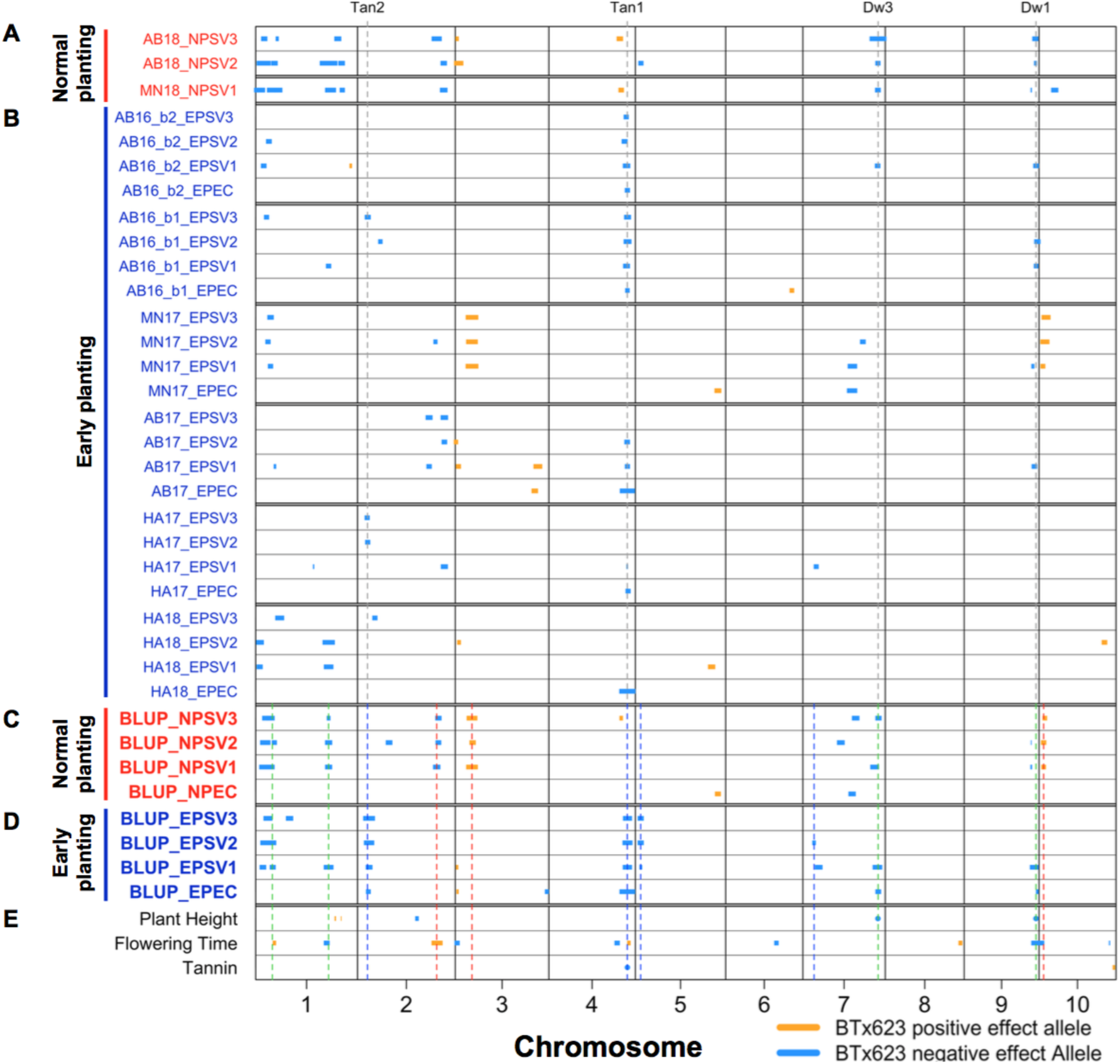
Joint linkage mapping (JLM) of chilling tolerance and undesirable traits. JLM of seedling traits from individual (A) normal (NP) and (B) early planting (EP) field trials. Field location and year were included as prefixes for each seedling trait. Five NP traits that failed to detect QTL were excluded from the figure but used for calculating NP seedling trait BLUPs. JLM with seedling trait BLUPs, generated with ∼75,000 data points from ∼16,000 field plots, from (C) normal and (D) early planting. Additionally, (E) JLM of plant height, flowering time, and grain tannins were included. Classical dwarfing and tannin genes were noted with gray dashed lines. Chilling tolerance QTL detected under early planting are noted with blue dashed lines and green lines noted chilling tolerance QTL detected under both early and normal planting. The QTL under normal planting were noted with red dashed lines. Positive or negative effects of the BTx623 allele was indicated in orange or blue colors, respectively. The percentage of variation explained is proportional to the width of the box for each locus and loci explaining phenotypic variation >10% are noted with circles. Abbreviations: EC, emergence count. SV1–3, seedling vigor1–3.

To check the stability of QTL across locations and years, JLM was performed separately by location. The QTL on chromosome 9 was detected in three early-planted locations, while QTL on chromosomes 2 and 7 were mapped in two locations (Figure 3B and S13–S18). The chromosome 4 QTL was consistently detected across early-planted field locations and years. The only exception was the MN17 field trial, which emerged under optimal conditions and experienced chilling one week later, where the chromosome 4 QTL was not detected (Figures 3B and S16). Among the loci detected with JLM of field phenotypes from early-and normal-planted individual field trials ((Figures 3A–B), few overlaps were observed.

### Mapping for agronomic traits and grain tannin

CIM and JLM was conducted to identify loci underlying plant height, flowering time, and grain tannins. CIM detected three plant height QTL in the HKZa family (Table S6 and Figure S19), and two each in the NSZ and Kao families, explaining 30–82% of plant height variation. Two plant height QTL, detected on chromosomes 7 and 9, colocalized with classical dwarfing genes *Dw3* and *Dw1*, respectively (Multani *et al.* 2003; Hilley *et al.* 2016). JLM identified six plant height QTL, of which alleles at four and two QTL contained negative and positive effects, respectively (Figures 3C and S21, and Table 3). Three QTL of major effect explained 85% plant height variation. Major height loci were 12 kb and 0.1 Mb from *Dw3* and *Dw1* genes, respectively.

**Table 3.**
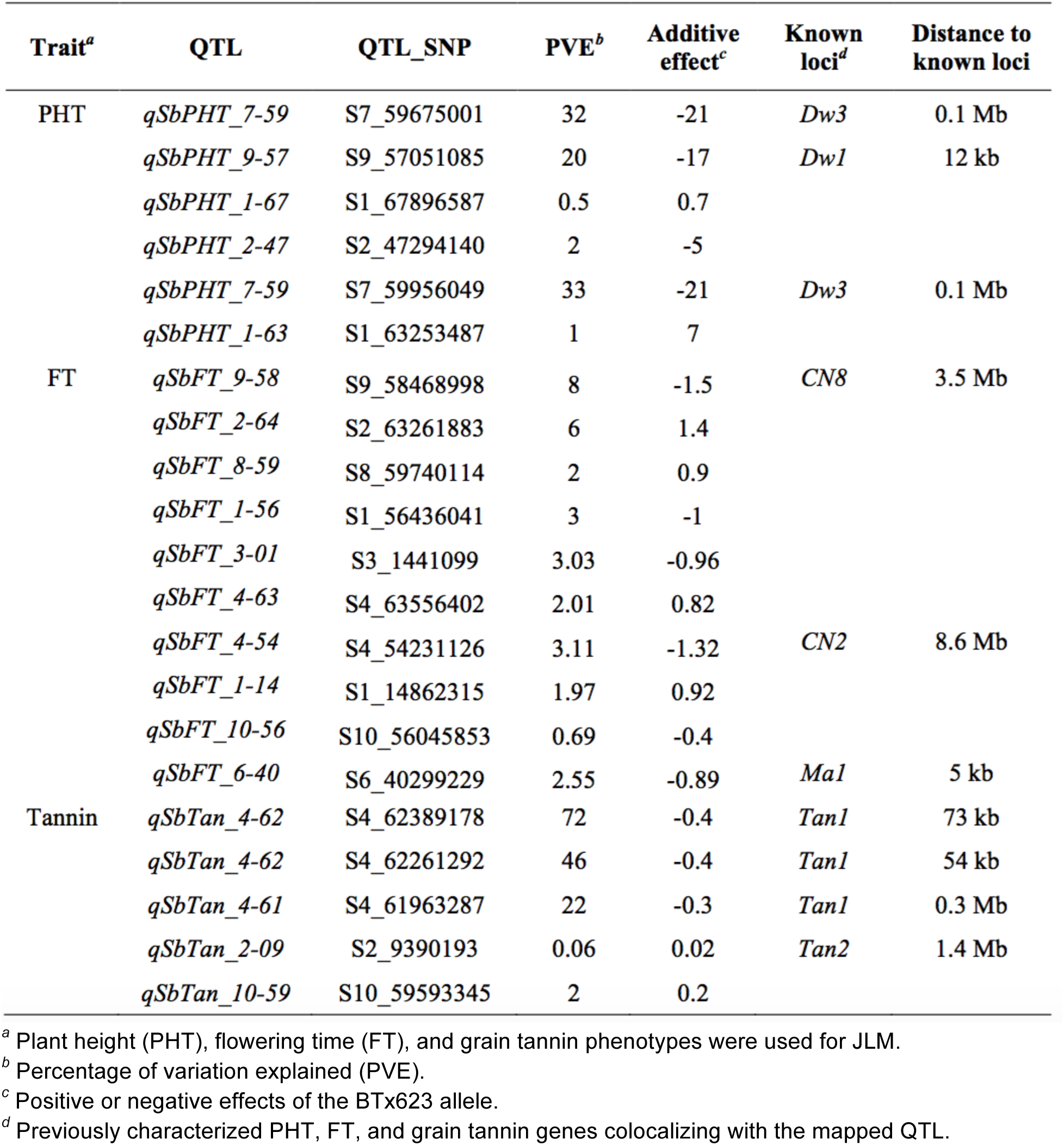
Joint linkage mapping of plant height, flowering time, and grain tannins.

| Trait<sup>a</sup> | QTL | QTL_SNP | PVE<sup>b</sup> | Additive effect<sup>c</sup> | Known loci<sup>d</sup> | Distance to known loci |
| --- | --- | --- | --- | --- | --- | --- |
| PHT | *qSbPHT_7-59* | S7_59675001 | 32 | -21 | *Dw3* | 0.1 Mb |
|  | *qSbPHT_9-57* | S9_57051085 | 20 | -17 | *Dw1* | 12 kb |
|  | *qSbPHT_1-67* | S1_67896587 | 0.5 | 0.7 |  |  |
|  | *qSbPHT_2-47* | S2_47294140 | 2 | -5 |  |  |
|  | *qSbPHT_7-59* | S7_59956049 | 33 | -21 | *Dw3* | 0.1 Mb |
|  | *qSbPHT_1-63* | S1_63253487 | 1 | 7 |  |  |
| FT | *qSbFT_9-58* | S9_58468998 | 8 | -1.5 | *CN8* | 3.5 Mb |
|  | *qSbFT_2-64* | S2_63261883 | 6 | 1.4 |  |  |
|  | *qSbFT_8-59* | S8_59740114 | 2 | 0.9 |  |  |
|  | *qSbFT_1-56* | S1_56436041 | 3 | -1 |  |  |
|  | *qSbFT_3-01* | S3_1441099 | 3.03 | -0.96 |  |  |
|  | *qSbFT_4-63* | S4_63556402 | 2.01 | 0.82 |  |  |
|  | *qSbFT_4-54* | S4_54231126 | 3.11 | -1.32 | *CN2* | 8.6 Mb |
|  | *qSbFT_1-14* | S1_14862315 | 1.97 | 0.92 |  |  |
|  | *qSbFT_10-56* | S10_56045853 | 0.69 | -0.4 |  |  |
|  | *qSbFT_6-40* | S6_40299229 | 2.55 | -0.89 | *Ma1* | 5 kb |
| Tannin | *qSbTan_4-62* | S4_62389178 | 72 | -0.4 | *Tan1* | 73 kb |
|  | *qSbTan_4-62* | S4_62261292 | 46 | -0.4 | *Tan1* | 54 kb |
|  | *qSbTan_4-61* | S4_61963287 | 22 | -0.3 | *Tan1* | 0.3 Mb |
|  | *qSbTan_2-09* | S2_9390193 | 0.06 | 0.02 | *Tan2* | 1.4 Mb |
|  | *qSbTan_10-59* | S10_59593345 | 2 | 0.2 |  |  |

<sup>a</sup> Plant height (PHT), flowering time (FT), and grain tannin phenotypes were used for JLM.
<sup>b</sup> Percentage of variation explained (PVE).
<sup>c</sup> Positive or negative effects of the BTx623 allele.
<sup>d</sup> Previously characterized PHT, FT, and grain tannin genes colocalizing with the mapped QTL.

Although flowering time varied little among the founders (Figure 1E), transgressive segregation enabled detection of seven flowering time loci (four, two, and one QTL in the NSZ, Kao, and HKZa families, respectively) which explain 20–28% of variation (Table S6). JLM with flowering time detected 10 QTL that explained 33% variation (Figures 3C and S21, and Table 3), three of which co-localized with previously identified flowering time/maturity genes, *TOC1*/*CN2, ma1*, and *CN8*. CIM of grain tannin presence/absence identified a major QTL on chromosome 4 in each family, with the Chinese parent conferring tannin presence allele in each case (Figure S20). The locus colocalizing with *Tan1* explained 77, 34, and 100% of grain tannin variation in the HKZa, NSZ, and Kao families, respectively (Table S6). JLM identified two tannin loci, one mapped ∼70 kb from *Tan1* and the other mapped ∼1.4 Mb from an earlier reported *Tan2* candidate gene (Wu *et al.* 2012; Morris *et al.* 2013b) (Figure S22, and Table 3).

## Discussion

### A NAM resource to dissect the genetic architecture of chilling tolerance

Characterizing the genetic architecture of adaptive traits provides insight into mechanisms of adaptation (Orr 2005) and guides strategies for breeding (Bernardo 2008). The NAM approach has been used to increase power and accuracy for dissection of complex adaptive traits in several widely adapted crop species (Buckler *et al.* 2009; Nice *et al.* 2016; Bouchet *et al.* 2017). By using temperate-adapted founders with contrasting chilling responses (Figures 1C, 1D, and S4), the chilling NAM resource addresses a gap in available sorghum NAM resources (Bouchet *et al.* 2017). Together, the chilling NAM and global NAM population (Bouchet *et al.* 2017) make up a resource of >3000 lines for complex trait dissection in sorghum. Given the founder lines originated from different botanical races (kafir-caudatum vs. durra; Figure 2A), the chilling NAM population should harbor abundant diversity for future studies of adaptive traits. Anecdotal field observations suggest the population harbors variation in vegetative pigmentation, disease susceptibility, and panicle and stem architecture.

The quality of the chilling NAM resource (i.e. RILs and corresponding SNP genotypes) developed in our study is validated by the precise mapping (<100 kb) of cloned dwarfing (*Dw1* and *Dw3*) and tannin (*Tan1*) genes (Figure 3, Table 3). Similarly, several major QTL (*qSbCT04.62, qSbCT02.08, qSbCT07.59*, and *qSbCT09.57*) were encompassed within the QTL intervals detected previously (Knoll *et al.* 2008; Burow *et al.* 2010) (Table S7). Notably, however, the greater population size (∼4–5-fold) and marker density (>100-fold) with NAM relative to earlier studies greatly improved the mapping resolution (>10-fold; Table S7) and power (i.e. several additional loci identified). Family structure and LD decay of the chilling NAM population generally matches expectations based on population design and observations from previous NAM populations (Bouchet *et al.* 2017). Genotypic (Figure 2A) and phenotypic similarity of HKZa and HKZb RILs suggest that the differentiation is due to residual heterozygosity in the HKZ founder or pollen contamination from another Chinese accession. Uncertainty regarding the pedigree of HKZb RILs does not diminish their usefulness as a part of the NAM resource (e.g. Figure 3).

QTL mapping from multi-environment trials clearly identified a major oligogenic component of chilling tolerance (Figure 3), consistent with previous work (Knoll *et al.* 2008; Burow *et al.* 2010; Fiedler *et al.* 2016; Ortiz *et al.* 2017). In keeping with the breeding goals, we considered all QTL that controlled performance under chilling stress (emergence, seedling vigor, or both) as chilling tolerance loci (Table 2), regardless of whether they also controlled performance under normal conditions. As chilling tolerance trials were conducted in a field environment, heritability and QTL effect sizes (Tables 1 and 2) were somewhat reduced compared to previous experiments under controlled conditions (Knoll *et al.* 2008). While replicability of field phenotyping for abiotic stress is a major challenge (Araus and Cairns 2014), observing plant performance under field conditions may increase the likelihood that genetic discoveries will translate to farmer fields (Cobb *et al.* 2018). A common limitation for molecular breeding of stress tolerance has been a lack of QTL stability (i.e. QTL × environment interaction) (Bernardo 2008). The overlapping of multi-environment chilling tolerance QTL from this study with QTL previously identified in the fields in Texas and Indiana (Table S7) provides evidence of their stability across a wide range of early-season chilling scenarios.

### The genetic basis of early-season chilling tolerance

Molecular networks for cold sensing and response appear to be largely conserved across plants (Knight and Knight 2012; Dong *et al.* 2019). These findings are consistent with long-standing observations of homologous variation in cold tolerance across diverse grasses, including sorghum (Vavilov 1951). For this reason, we considered whether NAM provides evidence that chilling tolerance in Chinese sorghum is due to derived variation at canonical cold tolerance genes (e.g. *CBF*s, *COLD1, SENSITIVE TO FREEZING2*, etc). Overall, we found little evidence that the chilling tolerance in Chinese sorghum is due to variation in canonical cold regulators (i.e. little localization between QTL and sorghum orthologs of known plant cold tolerance genes). The most significant and consistent QTL (*qSbCT04.62*; Table 2) colocalized with CBF gene Sobic.004G283201 (120 kb from the peak SNP), ortholog of the canonical Arabidopsis cold acclimation regulator *CBF1* (Thomashow 2001; Park *et al.* 2015). However, the sequence of the CBF gene from the Chinese founders revealed no change in their predicted peptide, and a previous study showed no chilling-responsive expression in chilling-tolerant NSZ (Marla *et al.* 2017). These findings suggest that a different closely linked gene, or the nearby *Tan1* gene, underlie this chilling tolerance QTL. No other QTL colocalized with orthologs of known plant cold tolerance genes (Thomashow 2001; Welti *et al.* 2002; Moellering *et al.* 2010).

The chilling tolerance QTL observed in our study may represent novel chilling tolerance mechanisms in sorghum, or conserved mechanisms not yet described in model plants. Fine-mapping and positional cloning of each chilling tolerance QTL (Ma *et al.* 2015) will be needed to address these or other hypotheses on the molecular basis of chilling tolerance in sorghum. Still, the genetic architecture provides some potential clues. Surprisingly, chilling tolerance QTL colocalized closely with classical tannin (*Tan1* and *Tan2*) and dwarfing genes (*Dw1* and *Dw3*) (Figure 3), four of the five most important genes under selection by US sorghum breeders in the 20th century (the fifth important gene, not colocalizing with chilling tolerance QTL is *Maturity1*) (Karper and Quinby 1946; Stephens *et al.* 1967; Wu *et al.* 2012; Morris *et al.* 2013a). This finding contradicted our original hypothesis of weak coupling-phase linkage of chilling susceptibility alleles with nontannin and dwarfing alleles. The colocalization itself could be due to (i) tight linkage (e.g. <1 Mb) of chilling tolerance loci to classical tannin and dwarfing loci or (ii) pleiotropic effects of classical tannin and dwarfing loci on chilling tolerance.

First we considered whether coinheritance of tannin and chilling tolerance alleles could be due to a pleiotropic effect of seed pigmentation regulators (*Tan1* and *Tan2*) on chilling tolerance. A conserved MBW ternary complex controls biosynthesis of flavonoids and tannins in plants via interactions of Myb and bHLH transcription factors with a WD40 transcriptional regulator (Nesi *et al.* 2000; Gu *et al.* 2011; Gao *et al.* 2018). Among sorghum tannin genes, *Tan1* encodes the WD40 component (Wu *et al.* 2012) and *Tan2* colocalizes with the bHLH transcription factor (Sobic.002G076600) (Morris *et al.* 2013b) orthologous to Arabidopsis *TRANSPARENT TESTA8* (*AtTT8*) and rice red grain gene (*OsRc*) (Nesi *et al.* 2000; Gu *et al.* 2011). The MBW complex has pleiotropic effects on abscisic acid-mediated seed dormancy and polyphenol-mediated protection from soil-borne pathogens (Helsper *et al.* 1994; Gu *et al.* 2011; Jia *et al.* 2012), which could contribute to emergence and seedling vigor under chilling. The chilling tolerance QTL *qSbCT02.08* detected in JLM of nontannin RILs (Figure S23) suggests that early-season chilling tolerance does not require seed tannins, even if the trait is under the control of the MBW complex. The existence of a Chinese accession Gai Gaoliang (PI 610727) that is chilling-tolerant but lacks grain tannins (Burow *et al.* 2010) supports this hypothesis.

Next we considered whether plant height alleles (*Dw1* and *Dw3*) could have pleiotropic effects on chilling tolerance that explain their colocalization with *qSbCT07.59* and *qSbCT09.57* (Figure S9). *Dw1*, which colocalized with *qSbCT09.57*, encodes a novel component of brassinosteroid (BR) signaling (Hirano *et al.* 2017). BR signaling controls cold tolerance mechanisms in tomato (Xia *et al.* 2018) and Arabidopsis (Eremina *et al.* 2016) so colocalization of *qSbCT09.57* with *Dw1* could reflect a pleiotropic chilling tolerance effect of DW1 BR signaling. *Dw3*, which colocalized with *qSbCT07.59*, encodes an auxin transporter. However, to our knowledge, no reports have demonstrated a role of auxin signaling in chilling tolerance.

### Origins and consequences of the genetic architecture of chilling tolerance

Chilling sensitivity of US sorghum has generally been understood to be a result of sorghum’s tropical origin (Stickler *et al.* 1962; Knoll *et al.* 2008) (Figure 4A), in keeping with a classic phytogeographic model (Vavilov 1951). Under this model, ancestrally chilling-sensitive African sorghums would have adapted to cold upon diffusion to temperate regions in central Asia and northern China (c. 800 years ago) due to derived alleles (Kimber 2000). However, our finding that chilling tolerance alleles coinherited with the ancestral wildtype alleles of classical tannin and dwarfing genes, which are widespread in both African and Chinese sorghums, contradicts this original model.

**Figure 4.**
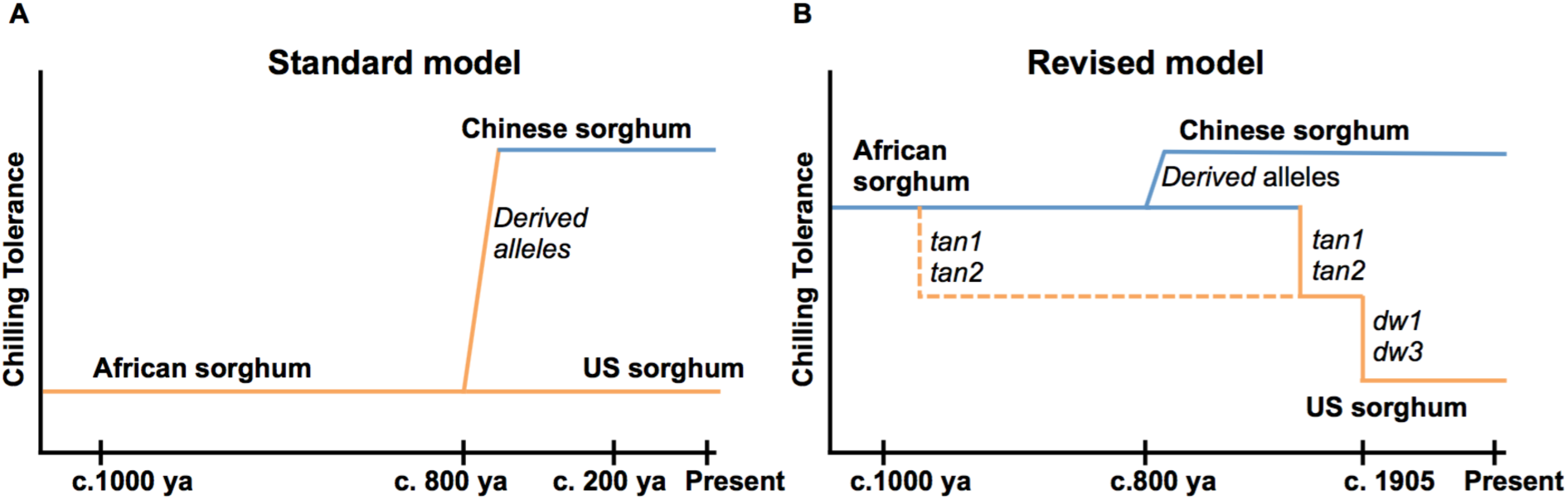
Evolutionary origin and agronomic effects of chilling tolerance. (A) Standard model: African sorghums are chilling sensitive based on their tropical origin, sorghum dispersed into northern China (c. 800 years ago) has adapted to chilling while the US sorghums derived from African sorghums remain chilling-sensitive. (B) Based on the genetic architecture of early-season chilling tolerance, we revised the model to explain chilling sensitivity of US sorghums. Coinheritance of chilling tolerance loci with wildtype alleles of classical dwarfing (*Dw1* and *Dw3*) and tannin (*Tan1* and *Tan2*) genes suggest tropical-origin sorghums are chilling-tolerant. Inadvertent selection of chilling-sensitive alleles with favorable dwarfing (*dw1* and *dw3*) and nontannin (*tan1* and *tan2*) alleles resulted in persistence of chilling sensitivity in US sorghums, despite breeding for chilling tolerance over the past 50 years.

Instead, a revised model for derived chilling sensitivity of US sorghum and inadvertent selection may be more parsimonious (Figure 4B). Under this model, the African sorghums introduced into the US harbored basal chilling tolerance, but chilling sensitivity was inadvertently selected along with loss-of-function alleles at *tan1* and *tan2* (from African standing variation), and *dw1* and *dw3* (from *de novo* mutations in US) (Multani *et al.* 2003; Morris *et al.* 2013b; Hilley *et al.* 2016). Supporting this revised model, 38 RILs selected for agronomic suitability by the sorghum breeder (R.P.) were fixed for the chilling-susceptibility alleles (at *qSbCT09.57* and *qSbCT07.59*) that are coinherited with desired *dw1* and *dw3* alleles, respectively (Table S8). Thus, coinheritance of chilling susceptibility with desired traits likely stymied >50 years of chilling tolerance breeding in this crop (Stickler *et al.* 1962; Tiryaki and Andrews 2001; Yu and Tuinstra 2001; Knoll and Ejeta 2008; Burow *et al.* 2010; Kapanigowda *et al.* 2013).

A genotype-to-phenotype modeling approach, which couples genetic and ecophysiological modeling, can help assess the potential value of genotypes in a crop’s target population of environments (Cooper *et al.* 2014). Preliminary ecophysiological modeling suggests that (were it not for chilling sensitivity) a standard grain sorghum hybrid could escape drought and have higher yields (∼5%) if planted 30–60 days early (Figure S24). The improved power and resolution with the chilling NAM provides several new paths to obtain chilling tolerance while bypassing undesirable characteristics from Chinese sorghum. Several chilling tolerance alleles (at *qSbCT05.04, qSbCT07.10, qSbCT01.13*, and q*SbCT01.57*) are not coinherited with undesirable alleles for tannins and height (Figure 3) and can be used directly in marker-assisted introgression. Complementary dominance of *Tan1* and non-functional *tan2* (Wu *et al.* 2012) can be exploited to develop chilling-tolerant sorghums that retain the nontannin phenotype. If the standard model is correct (Figure 4A), rare recombinants identified with high-density markers will decouple chilling tolerance alleles from undesirable wildtype alleles of tannin and dwarfing genes and bypass undesirable coinheritance. If the revised model is correct (Figure 4B), antagonistic pleiotropic effects could be bypassed with novel tannin biosynthesis mutations to disrupt tannin production in *Tan1Tan2* chilling-tolerant background and novel dwarfing mutants (Jiao *et al.* 2016) in *Dw1Dw3* chilling-tolerant background.

### Conclusions

Genetic tradeoffs caused by linkage drag have long been appreciated by geneticists and breeders (Zhu *et al.* 2018; Cobb *et al.* 2018). More recently, genetic tradeoffs due to antagonistic pleiotropy or conditional neutrality (Anderson *et al.* 2011) have been revealed by positional cloning of key agronomic genes (i.e. those under strong selection in 20th century breeding programs). For instance, antagonistic pleiotropic effects were identified for key improvement alleles of rice *semi-dwarf1* (Li *et al.* 2018) and tomato *jointless* (Soyk *et al.* 2017). In elite rice germplasm, conditional neutrality led to unintentional fixation of a drought-susceptibility allele at *Deeper rooting1* (Uga *et al.* 2013). Similarly, our findings suggest that strong selection for nontannin alleles (*tan1* and *tan2*) and dwarfing alleles (*dw1* and *dw3*) in grain sorghum in the 20th century inadvertently resulted in the loss of early-season chilling tolerance, due either (i) to tight repulsion-phase linkage of desired alleles (Figure 4A) or (ii) antagonistic pleiotropic effects of desired alleles on chilling susceptibility (Figure 4B). Given increasing evidence of genetic tradeoffs for genes under strong directional selection, characterizing both the genetic architecture and molecular basis of adaptive variation will be critical to guide genomics-enabled breeding and understand adaptive mechanisms.

## Author Contributions

The study was conceived by G.B. and G.M. The population was developed by G.B, R.C, and C.H. Data collection was by S.M., T.F., and R.P. Data was analyzed by S.M., T.F., Z.H., and M.O. Crop simulations were done by R.R. The paper was written by S.M. and G.M. All authors edited and approved the manuscript.

## Acknowledgements

The authors would like to thank Halee Hughes and Matt Davis for excellent technical support. Development of the NAM was supported by USDA ARS CRIS#3096-21000-021-00D and United Sorghum Checkoff Program (USCP) Grant on “Sorghum Genetic Enhancement” to USDA-ARS, Lubbock, TX. Dr. Ratan Chopra was supported by the grant from United Sorghum Checkoff Program. The study was supported by the Kansas Grain Sorghum Commission and Kansas Department of Agriculture. The study was carried out using the Beocat high-performance computing facility and Integrated Genomics Facility at Kansas State University. This study is contribution no. [*add after acceptance*] from the Kansas Agricultural Experiment Station.

